# BEdeepoff: an *in silico* tool for off-target prediction of ABE and CBE base editors

**DOI:** 10.1101/2021.03.14.435296

**Authors:** Chengdong Zhang, Daqi Wang, Tao Qi, Yuening Zhang, Linghui Hou, Feng Lan, Jingcheng Yang, Sang-Ging Ong, Hongyan Wang, Leming Shi, Yongming Wang

## Abstract

Base editors, including adenine base editors (ABEs) and cytosine base editors (CBEs), are valuable tools for introducing point mutations, but they frequently induce unwanted off-target mutations. Here, we performed a high-throughput gRNA-target library screening to measure editing efficiencies at integrated genomic off-targets and obtained datasets of 48,632 and 52,429 off-targets for ABE and CBE, respectively. We used the datasets to train deep learning models, resulting in ABEdeepoff and CBEdeepoff which can predict editing efficiencies at off-targets. These tools are freely accessible via online web server http://www.deephf.com/#/bedeep.

## Introduction

Base editors enable programmable conversion of a single nucleotide in the mammalian genome and have a broad range of research and medical applications. They are fusion proteins that include a catalytically impaired Cas9 nuclease (Cas9^D10A^) and a nucleobase deaminase ^1, 2^. Cas9^D10A^ nuclease and a ~100 nucleotide guide RNA (gRNA) forms a Cas9^D10A^-gRNA complex, recognizing a 20 nucleotide target sequence followed by a downstream protospacer adjacent motif (PAM) ^3–5^. Once the Cas9^D10A^-gRNA complex binds to target DNA, it opens a single-stranded DNA loop ^3^. The nucleobase deaminase modifies the single-stranded DNA within a small ~5 nucleotide window at the 5’ end of the target sequence ^1, 2^. Two classes of base editors have been developed to date: cytidine base editors (CBEs) convert target C:G base pairs to T:A ^1^, and adenine base editors (ABEs) convert A:T to G:C ^2^. Base editors have been successfully used in diverse organisms including prokaryotes, plants, fish, frogs, mammals, and human embryos ^6–9^.

As a major concern of base editing, off-target editing can occur on genomic sequences that are similar to the 20-nucleotide target sequence ^10, 11^. Experimental evaluation of a target sequence is time-consuming, prompting us to develop *in silico* tools for target sequence evaluation. We recently used a high-throughput strategy for gRNA-target library screening for SpCas9 activity ^12^. In this study, we designed libraries of gRNA-off-target sequence pairs and performed high-throughput screen, obtaining 48,632 and 52,429 valid off-targets for ABE and CBE, respectively. The resulting datasets were used to train deep learning models, resulting in ABEdeepoff and CBEdeepoff which can predict editing efficiency at potential off-targets for ABE and CBE, respectively.

## Results

### A guide RNA-target pair strategy for test of editing efficiency at off-targets

To investigate editing efficiency at off-targets, we designed two gRNA-target pair libraries with one for ABE and one for CBE (Fig. 1a). Each library contains 91,566 gRNA-target pairs, distributed among 1,383 gRNA groups for ABE and 1,378 gRNA groups for CBE. Each group contains one gRNA-on-target pair and multiple gRNA-off-target pairs. The mutation type included mismatches (1-6 bp mismatches per off-target, 71,099 for ABE and 70,594 for CBE), deletions (1-2 bp per target, 6,521 for ABE and 6,759 for CBE), insertions (1-2 bp per target, 11,396 for ABE and 11,562 for CBE) and mismatches mixed with insertions/deletions (indels, 1-2 bp mismatches plus 1-2 bp indels, total mutant nucleotides are 2-3 bp, 888 for ABE and 881 for CBE). We also designed non-target sequences as control (279 for ABE and 392 for CBE).

**Figure 1.**
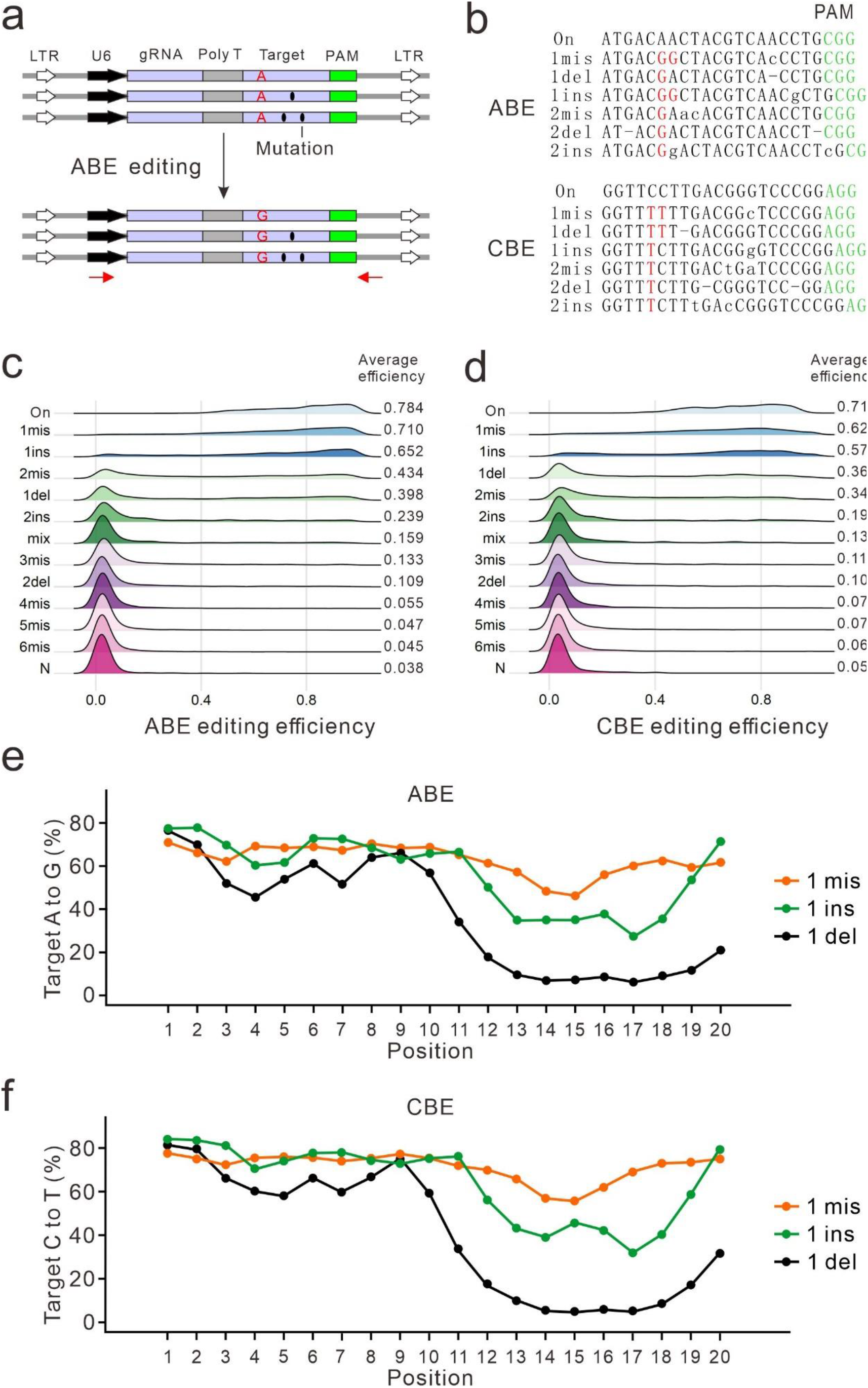
High-throughput screen of conversion efficiencies at off-targets. (**a**) A gRNA group of pairwise library for off-target screening. Black ellipses indicate mutations; red arrows indicate primers for target site amplification. (**b**) Deep-sequencing results reveal that A to G conversion for ABE and C to T conversion (red letter) for CBE occurred. Mismatched nucleotides and inserted nucleotides are indicated by lowercase letters; deletion is indicated by On: on-target sequence; 1mis: 1bp mismatch; 1del: 1 bp deletion; 1ins: 1 bp insertion. (**c, d**) Effects of the mutation type on nucleotide conversion efficiency. Mix: mismatches mixed with indels; On: on-target; N: 23 bp random sequence as negative control. (**e, f**) Positional effect on conversion efficiency for 1 bp mismatch, 1 bp insertion and 1 bp deletion.

To make the high-throughput data more reproducible, we first generated a panel of single cell-derived clones that stably express ABE or CBE base editors. Optimized version of base editors (ABEmax for ABE; AncBE4max for CBE) ^13^ were used in this study. Because of the large size, we integrated base editors into the genome by using the Sleeping Beauty (SB) transposon system (Supplementary Fig. 1a) ^14–16^. We tested conversion efficiency in each clone and selected an efficient clone for each base editor (Supplementary Fig. 1b-c).

We packaged the gRNA-target pair library into lentiviruses and transduced them into recipient cells. Five days after transduction, genomic DNA was extracted, and synthesized off-targets were PCR-amplified for deep-sequencing (Fig. 1a). Deep sequencing results revealed that A to G conversion for ABE and C to T conversion for CBE occurred (Fig. 1b). In this study, an off-target efficiency was defined as No. of edited reads divided by the No. of total valid reads. The screening assay was experimentally repeated twice, and editing efficiency in two independent replicates showed high correlation (Spearman correlation, 0.956 for ABE and 0.965 for CBE). Data from the two replicates were combined together for subsequent analysis. We obtained valid ABE efficiencies (reads number>100) of 48,632 for off-targets, 1,052 for on-targets and 153 for non-target controls. We obtained valid CBE efficiencies (reads number>100) of 52,429 for off-targets, 1,030 for on-targets and 220 for non-target controls.

The large-scale datasets generated here allowed us to analyze the effects of mutation type on editing efficiency. For both base editors, editing efficiencies decreased with an increasing number of mismatches (Fig. 1c-d). Deletions had a stronger influence than insertions and mismatches. The overall influence of mutation type on ABE could be ranked as 6mis > 5mis > 4mis > 2del > 3mis > mix > 2ins > 1del > 2mis > 1 ins > 1mis (Fig. 1c); the overall influence of mutation type on CBE could be ranked as 6mis > 5mis > 4mis > 2del > 3mis > mix > 2ins 2mis > 1del > 1ins > 1mis (Fig. 1d).

Next, we investigated the positional effects of mutation on editing efficiency. For both base editors, one mismatch only had minimal influence on editing efficiency at positions 14 and 15 but not at other positions; one insertion had minimal influence at the PAM-distal region but had stronger influence at positions 12-19; one deletion had mild influence at the PAM-distal region but had a stronger influence at PAM-proximal region (Fig. 1e-f). Interestingly, mutations at positions 19-20 had less influence than those at other seed region. Two mismatches had a stronger influence on editing efficiency when they both occurred on the seed region for both base editors (Supplementary Fig. 2a-b).

### Developing models for prediction of editing efficiency at off-targets

Next, we used ABE off-target editing datasets to train a shared embedding based deep learning model where a gRNA-off-target pair was considered as a unique sequence and used as input (Supplementary Fig. 3). The shared embedding design encoded the input gRNA and off-target sequences in the same scheme, and then embedded the representation vectors in the same matrix space. Both generalization ability and training speed will benefit from sharing the same embedding matrix instead of training an independent embedding matrix for each input. The gRNA-off-target pairs and non-target controls (48,632+153=48,785) were randomly split to two parts: one contained 41,906 for tenfold cross-validation with shuffling, and the other contained 4,879 for holdout testing (which were never seen by the model). The resulting model was named “ABEdeepoff” which can predict editing efficiency at a potential off-target. The model achieved high correlation for 1-2 bp mismatch, 1 bp insertion and 1 bp deletion (R>0.7), mild correlation for 2 bp insertions (R=0.57), weak correlation for 2 bp indels, 3 bp mismatches and mix mutations (R>0.3) and very weak correlation for the remaining mutations (R<0.03, Fig. 2a). These results suggest that ABEdeepoff performed well for off-targets with high editing efficiency but not with low editing efficiency.

**Figure 2.**
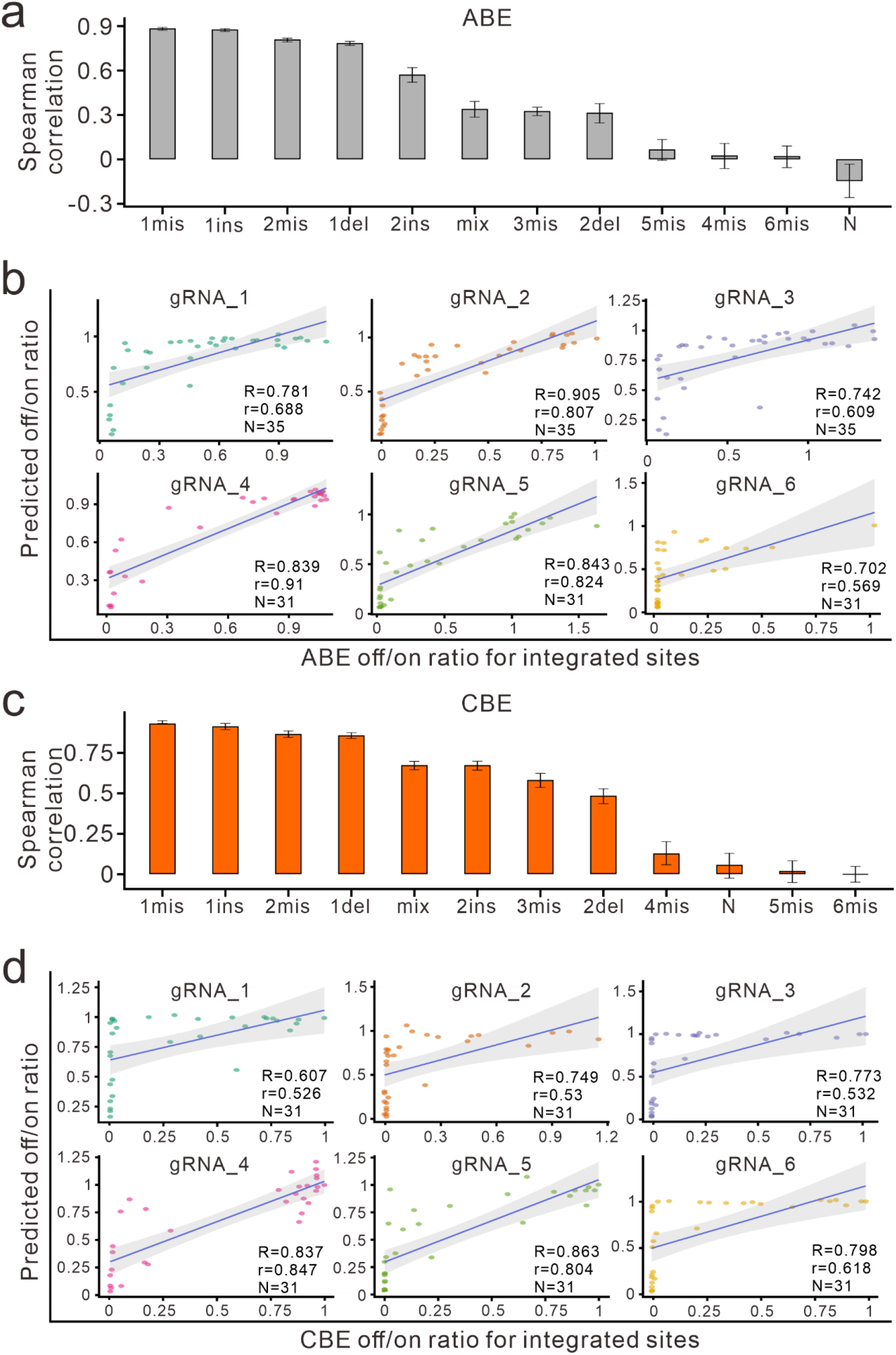
Evaluation of ABEdeepoff/CBEdeepoff for efficiency prediction at off-targets. (**a**) Evaluation of ABEdeepoff prediction for different mutation types with testing dataset. N: 23 bp random sequence as negative control. (**b)** Evaluation of ABEdeepoff prediction for conversion efficiency at six groups of integrated off-targets. (**c)** Evaluation of CBEdeepoff prediction for different mutation types with testing dataset. (**d)** Evaluation of CBEdeepoff prediction for conversion efficiency at six groups of integrated off-targets. R: Spearman correlation, r: Pearson correlation.

Next, we evaluated the performance of the model with six groups of integrated off-target datasets collected from literature ^10, 17^, and achieved Spearman correlation values that varied from 0.702 to 0.905 (Fig. 2b). Finally, we evaluated the performance of our model with 14 groups of endogenous off-target datasets collected from literature ^17^. We achieved a mild correlation (R>0.4) for seven targets and weak correlation for the remaining targets probably due to the low editing efficiencies measured at these off-targets (Supplementary Fig. 4).

Parallelly, we used the CBE off-target editing datasets to train a shared embedding based deep learning model. The gRNA-off-target pairs and non-target controls (52,429+220=52,649) were randomly split to two parts: one contained 47,384 for tenfold cross-validation with shuffling, and the other contained 5,265 for holdout testing. The resulting model was named “CBEdeepoff” which can predict editing efficiencies at potential off-targets. The model achieved high correlation (R>0.6) for 1-2 bp mismatch, 1-2 bp insertion, 1 bp deletion and mix mutations, mild correlation (R>0.4) for 3 bp mismatches and 2 bp deletions, and very weak correlation (R<0.2) for the remaining mutations (Fig. 2c). These results can be explained by that CBEdeepoff performed well for off-targets with high editing efficiency but not with low editing efficiency.

Next, we evaluated the performance of the model with six integrated groups of off-target datasets collected from literature ^11, 17^, and achieved Spearman correlation values varied from 0.607 to 0.863(Fig. 2d). Finally, we evaluated the performance of the model with seven groups of endogenous off-target datasets collected from literature ^17^. We achieved mild correlation (R>0.4) for four targets and weak correlation for the remaining targets probably due to the very low editing efficiencies measured at these off-targets (Supplementary Fig. 5).

ABEdeepoff /CBEdeepoff can be used together with Cas-OFFinder ^18^ and CRISPRitz ^19^. Both of the software can rapidly identify potential off-targets based on sequence similarity in the whole genome, and ABEdeepoff /CBEdeepoff can calculate editing efficiency for each sequence to define the potential off-targets (Fig. 3). The output file of Cas-OFFinder and CRISPRitz can be used as input file for ABEdeepoff or CBEdeepoff. We finally provided an online webserver named BEdeep, which is available at http://www.deephf.com/#/bedeep.

**Figure 3.**
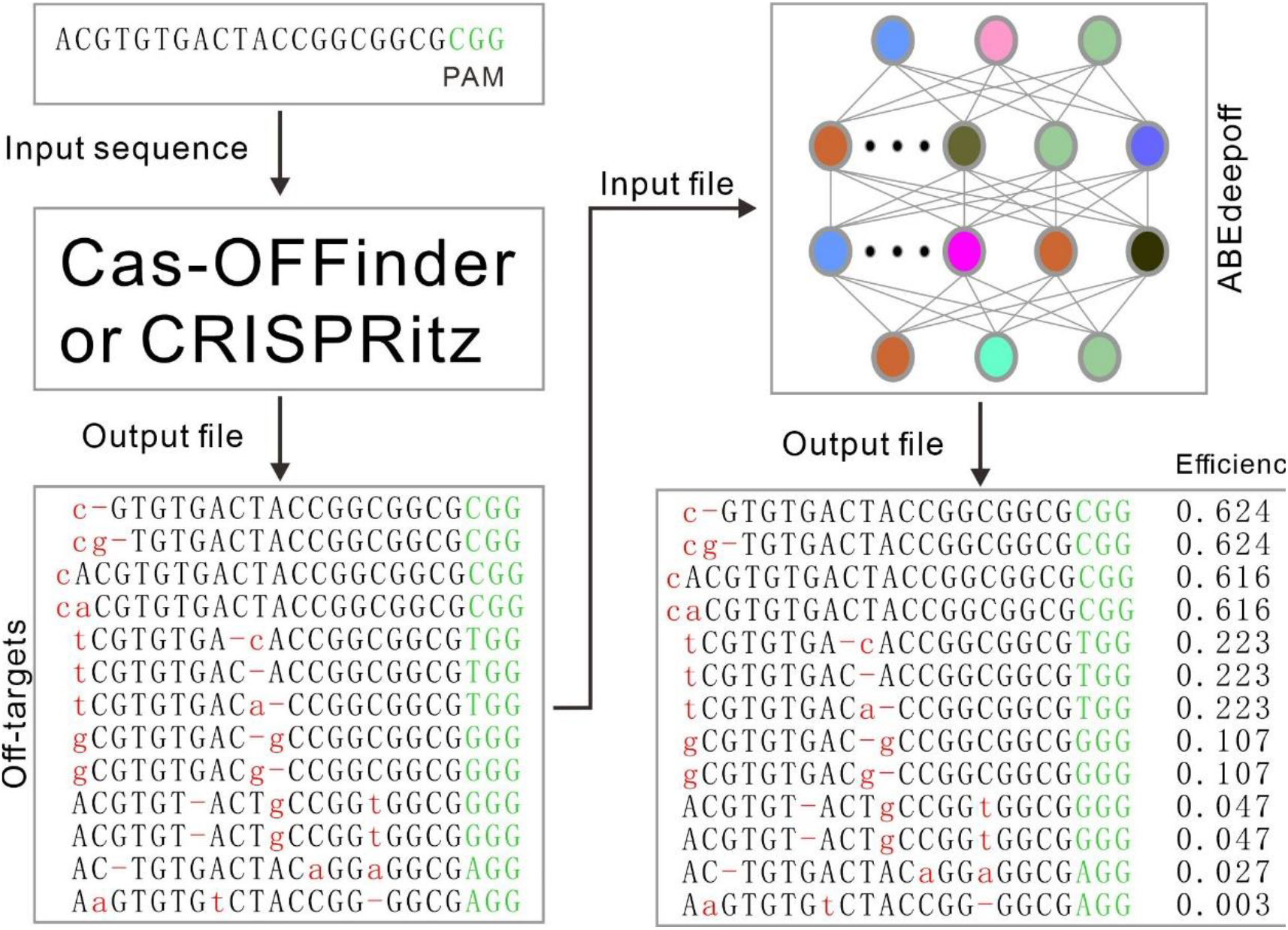
Example of ABEdeepoff for off-target identification. A target sequence is input into Cas-OFFinder or CRISPRitz to identify off-targets in human genome based on sequence similarity. Here we set up the parameters as maximal 2 mismatches, 1 deletion and 1 insertion. The output file containing potential off-targets is used as input file for ABEdeepoff. ABEdeepoff calculates editing efficiency for each potential off-targets.

## Discussion

Off-target editing is always a major concern of genome editing. As Cas9-deaminase fusion proteins, both ABE and CBE base editors can induce sequence independent and dependent off-target mutations ^10, 11, 20–22^. By engineering deaminases, sequence independent off-target mutations can be minimized to background level ^20, 23^. However, how to minimize sequence dependent off-target mutations is a challenge because eukaryote genomes contain hundreds of sequences similar to the target sequences. In this study, we developed models to predict editing efficiencies on potential off-targets for both ABE and CBE, facilitating users to select highly specific targets.

The shared embedding based deep learning models used here has a simple yet elegant structure which unifies on-target and off-target input in form. It can be seamlessly extended to other versions of base editors, such as SauriABEmax, SauriBE4max, SaKKH-BE3, BE4-CP, dCpf1-BE and eA3A-BE3 ^24–28^. It may also be used to generate models to predict editing outcomes for Cas9 and Cas12 nucleases.

## Methods

### Cell culture and transfection

HEK293T cells (ATCC) were maintained in Dulbecco’s Modified Eagle Medium (DMEM) supplemented with 10% FBS (Gibco) at 37 °C and 5% CO2. All media contained 100 U/ml penicillin and 100 mg/ml streptomycin. For transfection, HEK293T cells were plated into 6-well plates, DNA mixed with Lipofectamine 2000 (Life Technologies) in Opti-MEM according to the manufacturer’s instructions. Cells were tested negative for mycoplasma.

### Plasmid construction

SB transposon (pT2-SV40-BSD-ABEmax and pT2-SV40-BSD-BE4max) was constructed as follows: first, we replaced the Neo^R^ gene (AvrII-KpnI site) on pT2-SV40-Neo^R^ with BSD, resulting in pT2-SV40-BSD vector; second, backbone fragment of pT2-SV40-BSD was PCR-amplified with Gibson-SV40-F and Gibson-SV40-R, and ABEmax fragment was PCR-amplified from pCMV_ABEmax_P2A_GFP (Addgene#112101) with Gibson-ABE/BE4-F and Gibson-ABE/BE4-R, and BE4max fragment was PCR-amplified from pCMV_AncBE4max (Addgene#112094) with Gibson-ABE/BE4-F and Gibson-ABE/BE4-R; third, the backbone fragments were ligated with ABEmax and BE4max using Gibson Assembly (NEB), resulting in pT2-SV40-BSD-ABEmax and pT2-SV40-BSD-BE4max, respectively.

### Generation of cell lines expressing ABEmax or BE4max

HEK293T cells were seeded at ~40% confluency in a 6-well dish the day before transfection, 2 μg of SB transposon (pT2-SV40-BSD-ABEmax or pT2-SV40-BSD-BE4max) and 0.5 μg of pCMV-SB100x were transfected using 5 μl of Lipofectamine 2000 (Life Technologies). After 24 h, cells were selected with 10 μg/ml of blasticidin for 10 days. Single cells were sorted into 96-well plates for colony formation. Conversion efficiency was performed to screen cell clones with high levels of ABEmax and BE4max expression.

### The gRNA-off-target library construction

The off-target library was designed as follows: both ABEs and CBEs libraries contained 91,556 oligonucleotides, distributed among 1,383 and 1,378 gRNA groups for ABEs and CBEs, respectively; for ABEs, the off-target library included 51,170 1bp-mismatch, 10,889 2bp-mismatch, 10,310 1bp-insertion, 4,960 1bp-deletion, 2,261 3bp-mismatch, 2,264 4bp-mismatch, 2,255 5bp-mismatch, 2,260 6bp-mismatch, 1,086 2bp-insertion, 1,561 2bp-deletion, 888 mix, 279 non-target and 1,383 on-target; for CBEs, the off-target library included 50,676 1bp-mismatch, 11,054 2bp-mismatch, 10,481 1bp-insertion, 5,128 1bp-deletion, 2,221 3bp-mismatch, 2,226 4bp-mismatch, 2,203 5bp-mismatch, 2,214 6bp-mismatch, 1,081 2bp-insertion, 1,631 2bp-deletion, 881 mix, 392 non-target and 1,378 on-target; each oligonucleotide chain contained left sequence (tgtggaaaggacgaaacacc), gRNA sequence (NNNNNNNNNNNNNNNNNNNN), BsmBI site (gttttgagacg), Barcode 1 (NNNNNNNNNNNNNNNNNNNN), BsmBI site (cgtctcgctcc), Barcode 2 (NNNNNNNNNNNNNNN), target sequence (gtactNNNNNNNNNNNNNNNNNNNNNgg), and right sequence (cttggcgtaactagatct). The off-target library was constructed as follows: first, full-length oligonucleotides were PCR-amplified and cloned into BsmBI site of LentiGuide-U6-del-tracRNA vector by Gibson Assembly (NEB), named LentiGuide-U6-gRNA-target; second, the tracRNA were PCR-amplified and cloned into BsmBI site of LentiGuide-U6-del-tracrRNA vector by T4 DNA ligase (NEB).The Gibson Assembly products or T4 ligation products were electroporated into MegaX DH10B™ T1^R^ Electrocomp^TM^ Cells (Invitrogen) using a GenePulser (BioRad) and grown at 32 °C, 225 rpm for 16 h. The plasmid DNA was extracted from bacterial cells using Endotoxin-Free Plasmid Maxiprep (Qiagen).

### Lentivirus production

Lentivirus production was described previously^12^. Briefly, for individual sgRNA packaging, 1.2 μg of gRNA expressing plasmid, 0.9 μg of psPAX2 and 0.3 μg of pMD2.G (Addgene) were transfected into HEK293T cells by Lipofectamine 2000 (Life Technologies). Medium were changed 8 hours after transfection. After 48h, virus supernatants were collected. For library packaging, 12 μg of plasmid library, 9 μg of psPAX2, and 3 μg of pMD2.G (Addgene) were transfected into a 10-dish HEK293T cells with 60 μl of Lipofectamine 2000. Virus were harvested twice at 48 h and 72 h post-transfection. The virus was concentrated using PEG8000 (no. LV810A-1, SBI, Palo Alto, CA), dissolved in PBS and stored at −80 °C.

### Analysis of individual gRNA conversion efficiency

HEK293T cells expressing ABEmax or BE4max were seeded at ~40% confluency in a 24-well dish the day before infection. After the infection of gRNA-target paired lentivirus, genomic DNA was extracted at suitable time points using QuickExtract DNA Extraction Solution (Epicentre). We amplified the target sequence by PCR with Q5 High-Fidelity 2X Master Mix (NEB). PCR products were purified with Gel Extraction Kit (Qiagen) and Sanger-sequenced. According to “.ab1” sequencing file, the conversion efficiency was compared.

### Screening experiments in HEK293T

HEK293T cells expressing ABEmax or BE4max were plated into 15 cm dish at ~30% confluence. After 24 h, cells were infected with library with at least 1000-fold coverage of each gRNAs. After 24 h, the cells were cultured in the media supplemented with 2 μg/ml of puromycin for 5 days. Cells were harvested and the genomic DNA was isolated using Blood & Cell Culture DNA Kits (Qiagen). The integrated region containing the gRNA coding sequences and target sequences were PCR-amplified using Q5 High-Fidelity 2X Master Mix (NEB). We performed 60-70 PCR reactions using 10 μg of genomic DNA as template per reaction for deep sequencing analysis. The PCR conditions: 98 °C for 2 min, 25 cycles of 98 °C for 7 s, 67 °C for 15 s and 72 °C for 10 s, and the final extension, 72 °C for 2 min. The PCR products were mixed and purified using Gel Extraction Kit (Qiagen). The purified products were sequenced on Illumina HiSeq X by 150-bp paired-end sequencing.

### Data analysis

FASTQ raw sequencing reads were processed to identify gRNA-off-target editing activity. The nucleotides in a read with quality score < 10 was masked with a character “N”. Due to the integrated design strategy, we first separated a read to designed gRNA region, scaffold region, and target region to extract the corresponding sequence. The designed gRNA was then aligned to the reference gRNA library to mark the reads. The target sequence was compared to the designed gRNA to mark if the target was edited. We screened out gRNAs with a total valid reading of less than 100. Then, the efficiency for a specific gRNA-off-target pair can be calculated by the following formula:

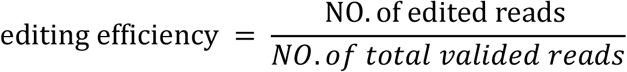

### Encoding

Drawing on concepts from the field of natural language processing (NLP), nucleotides A, C, G, and T can be regarded as words in a DNA sequences.

Therefore, we can widely use algorithms in the NLP field to solve prediction tasks in the CRISPR field, especially the use of embedding algorithms to get the continuous representation of discrete nucleotide sequences ^29^. Unlike the common efficiency prediction that only needs to input one single sequence for regression models, in this research, the gRNA-off-target pair has two different sequences as inputs. For gRNA, there are four words in the index vocabulary (i.e., A, C, G, and T); but after alignment, there might exist DNA bulge or RNA bulge and thus lead to a “-” word to represent the insertion or deletion in the gRNA or off-target sequence. So, the vocabulary can be described as:

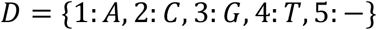

The input sequence can be described as:

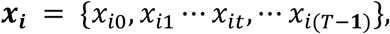

where *i* ∈{1,2}denotes the *i*-th sequence in an gRNA-off-target pair, *x_it_* is the t-th element of the *i*-th sequence, *T* is the sequence length. For example, ACGCTTCATCA-ATGTTGGGATGG (seq1, gRNA + NGG) and ACGC-TCATCAaAaGTT-GGATGG (seq2, off-target) can be encoded as:

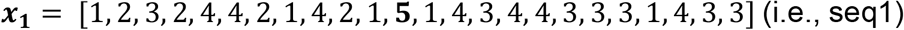

and

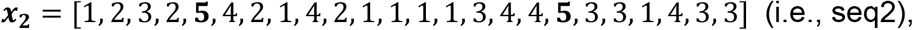

respectively.

### Shared embedding

Inspired by the algorithms in the recommender system ^30^ and click-through rate (CTR) ^31^ prediction modeling, both the generalization capacity and training speed will benefit from the sharing of the same embedding matrix instead of training independent embedding matrices for each input. In this research, a discrete nucleotide encoding *x_it_* is projected to the dense real-valued space 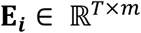 (*m* is a hyperparameter corresponds to the embedding dimension) to get the embedding vector **e**(*x_it_*). Then a final embedding matrix **E** is needed to get the combined information from those two embedding matrices by:

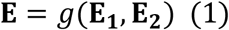

where *g* can be sum, mean, or even a simple concatenate function. However, the sum or mean function is more suitable because it can reduce the redundant features in **E_1_** and **E_2_**. We choose the sum function here for simplicity.

### Feature extraction and model prediction

Long-short-term memory network (LSTM) and gated recurrent unit network (GRU) are a type of recurrent neural networks (RNN) algorithms used to address the vanishing gradient problem in modelling time-dependent and sequential data tasks ^32^. Usually, a bidirectional manner was used to capture the information from the forward and backward directions of a sequence, which is biLSTM or biGRU. Our work and others’ work have shown that, as an important component, biLSTM can be used alone or with convolutional neural network (CNN) to achieve good performances in various regression and classification tasks involving biological sequences ^12, 33–35^. Here, we tried biLSTM, biGRU, and the newly proposed transformer structure ^36^, and found biLSTM had the fastest convergence speed. The input and output of biLSTM can be described by the following equations:

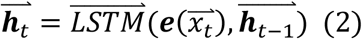

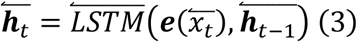

Thus, the output context vectors of biLSTM are 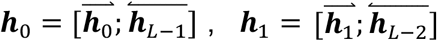, etc. Thus, we can concatenate the forward and backward hidden state as **H** = {***h***_0_, ***h***_1_,···, ***h***_*L*−1_}, which contains the bidirectional information in the shared embedding feature matrix. Before the fully connected layers, we tried different input features based on the trade-off of the convergence speed and the performance of the model. The aforementioned features are last hidden unit, max pooling operation on **H**, and average pooling on **H**. The equations are as following:

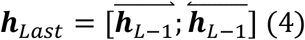

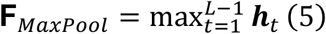

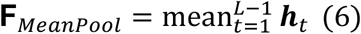

A max pooling and average pooling function were applied separately to **H** to obtain more useful features, and combine them with the final hidden state to produce the concatenated feature: ***c* =** [***h**_Last_*; **F**_*MaxPool*_; **F**_*MeanPool*_], the predicted efficiency *o* for a specific gRNA-off-target pair can be obtained by:

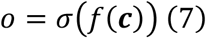

where, *f* is fully connected layers, *σ* is the *sigmoid* activation function. Considering that the sample sizes of off-target types vary greatly, in order to get better prediction results for the types with small sample sizes, we use the weighted MSE as the loss function.

### Training setting

Off-target datasets were randomly split into two parts, one for training, and the other for holdout testing. The stability of model performance was estimated by a 10-fold shuffled validation together with the external integrated and endogenous datasets. The ABEdeepoff and CBEdeepoff models share the following hyperparameters: embedding dimension, 256; LSTM hidden unites, 512; LSTM hidden layers, 1; dropout rate, 0.5; fully connected layers, 2 (6*256->3*256->1). For all the models, the Adam optimizer was used with a customized learning rate decay strategy that gradually reduce the learning rate from 0.001, 0.0001, 0.00005 to 0.00001.

### Tools used in the study

BWA-0.7.17 was used to identify the designed gRNA^37^. PyTorch 1.6 ^38^ was used for building deep learning models. The global algorithm of pairwise2 in BioPython 1.7.8 were used for getting the alignment result of off-target sequence.

## Data and code availability

The raw sequencing data have been submitted to the NCBI Sequence Read Archive (‘SRA PRJNA587328 [https://dataview.ncbi.nlm.nih.gov/object/19730669]’). The other data for the present study is available from the corresponding author upon reasonable request.

## Acknowledgments

This work was supported by grants from the National Natural Science Foundation of China (81870199, 81630087), the Foundation for Innovative Research Group of the National Natural Science Foundation of China (31521003), Science and Technology Research Program of Shanghai (Grant Number 19DZ2282100).

