## Supplemental figures for "BEdeepoff: an *in silico* tool for off-target prediction of ABE and CBE base editors"

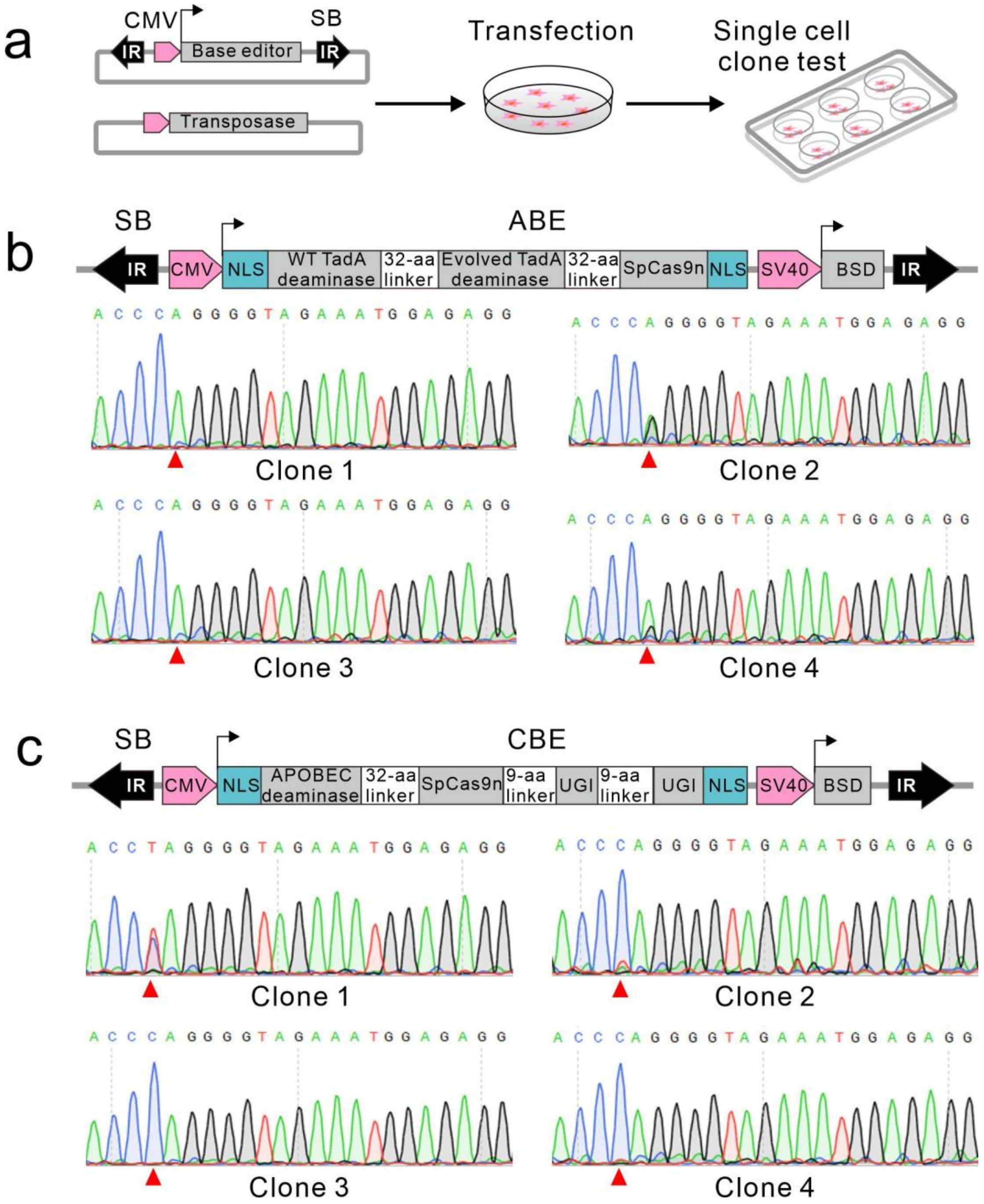


**Supplementary Figure 1. Generation of single cell-derived clones expressing base editors**

(**a**) Schematic diagram of cell clone generation. Sleeping Beauty (SB) transposon carrying base editors together with transposase vector are transfected into HEK293T cells followed by blasticidin selection. Five days after transfection, single cells are isolated and seeded into a new plate for colony formation. (**b, c**) Test of nucleotide conversion efficiency by editing EMX1 site in single cell-derived clones expressing ABEmax or CBEmax. Clone #2 is selected for ABE and Clone #1 is selected for CBE. Schematic diagrams of base editors are shown above. The edited positions are indicated by red triangles. IR: inverted repeat.

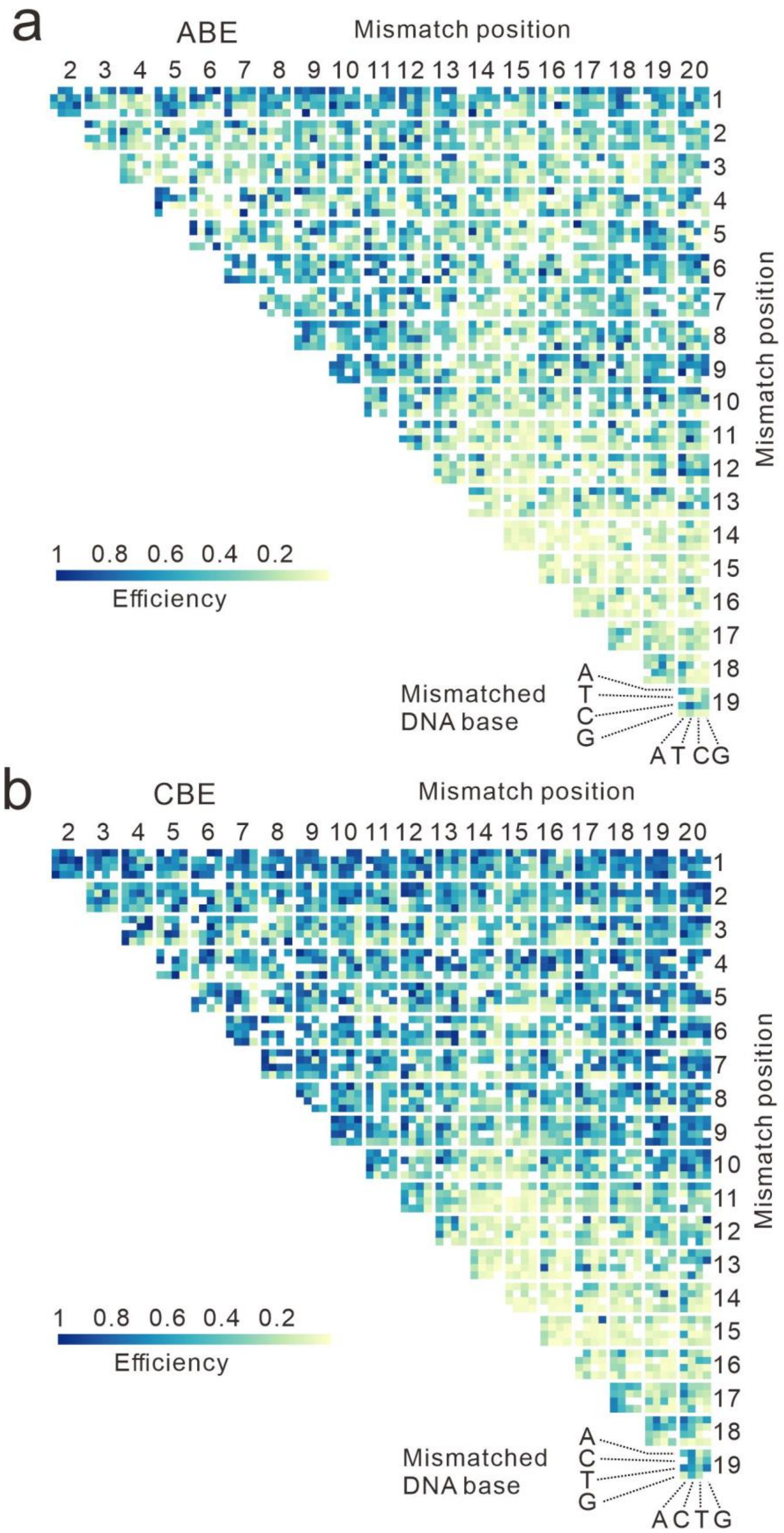


**Supplementary Figure 2. Positional effects of two nucleotide mismatches on conversion efficiency.**

(**a, b**) Influence of two nucleotide mismatches on conversion efficiency for ABE and CBE, respectively.

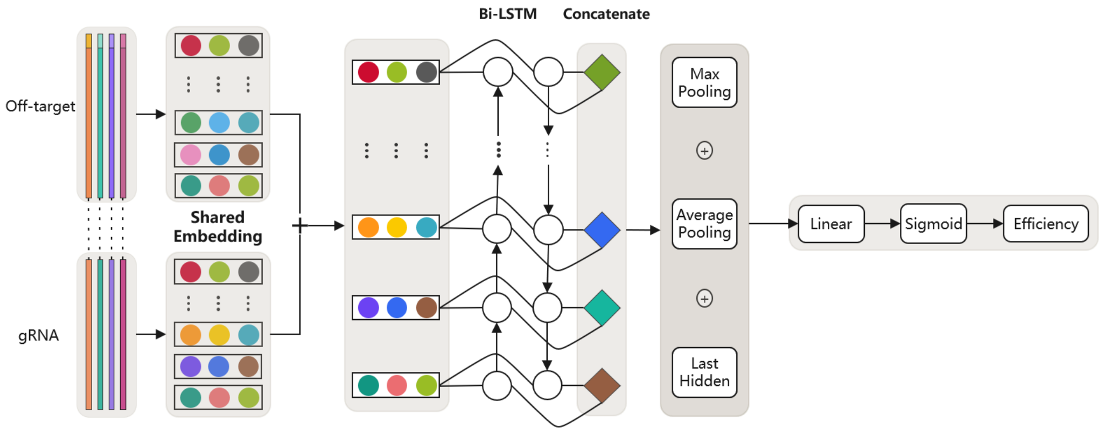


**Supplementary Figure 3. a shared embedding based deep learning model.** The gRNA and off-target sequence are paired together as input and embedded in the same matrix space to get the dense real-valued representation. Their representations are further processed by a matrix summation to get the combined features. This combined feature is further processed by a BiLSTM to get the final representation, which is then processed by the three feature extractors (i.e., Last Hidden, Average Pooling, and Max Pooling). The extracted features serve as the input of the fully connected layer. Finally, a sigmoid transformation is performed to get the predicted efficiency.

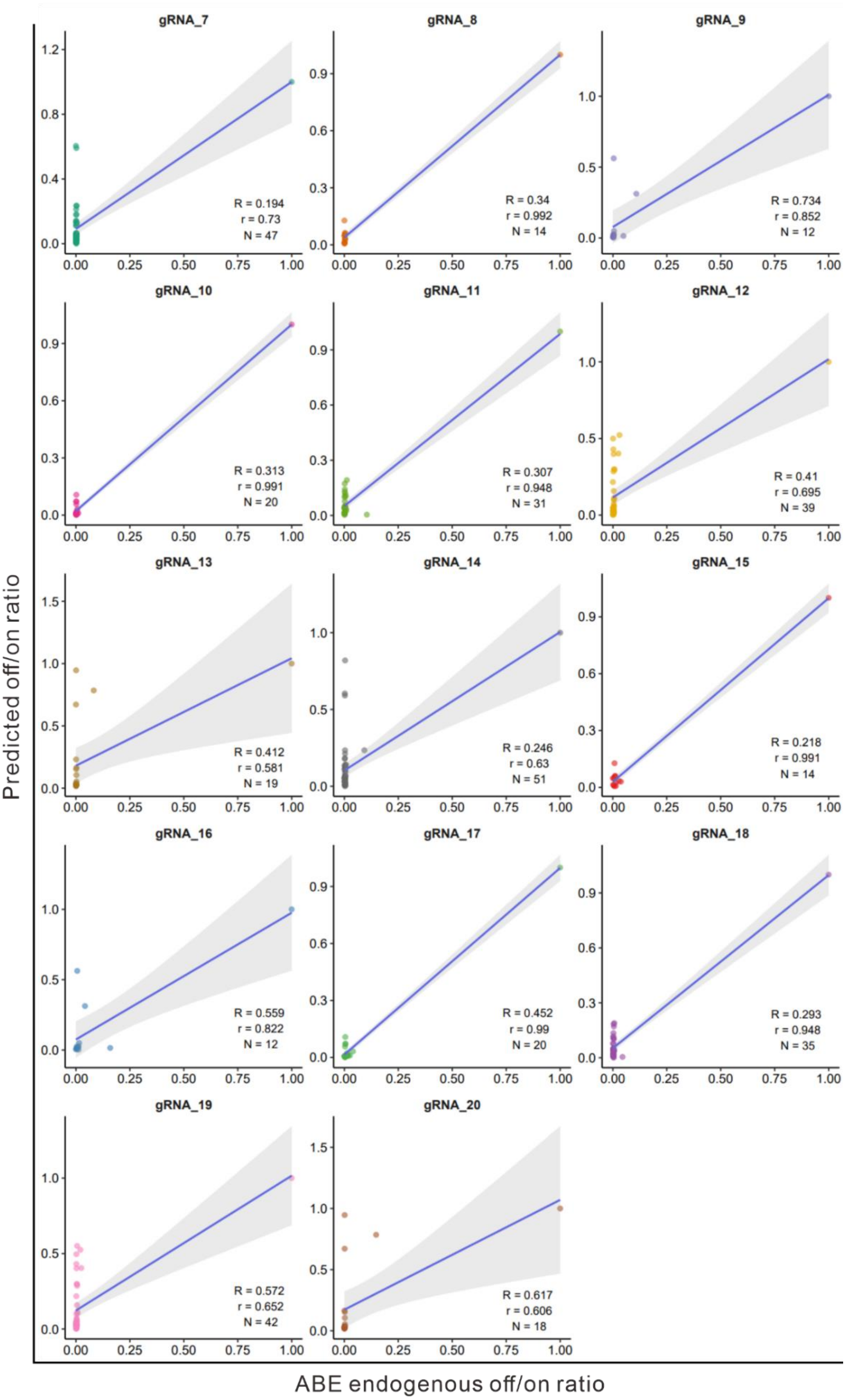


**Supplementary Figure 4. Evaluation of ABEdeepoff prediction for conversion efficiency with 14 groups of endogenous off-target datasets.**

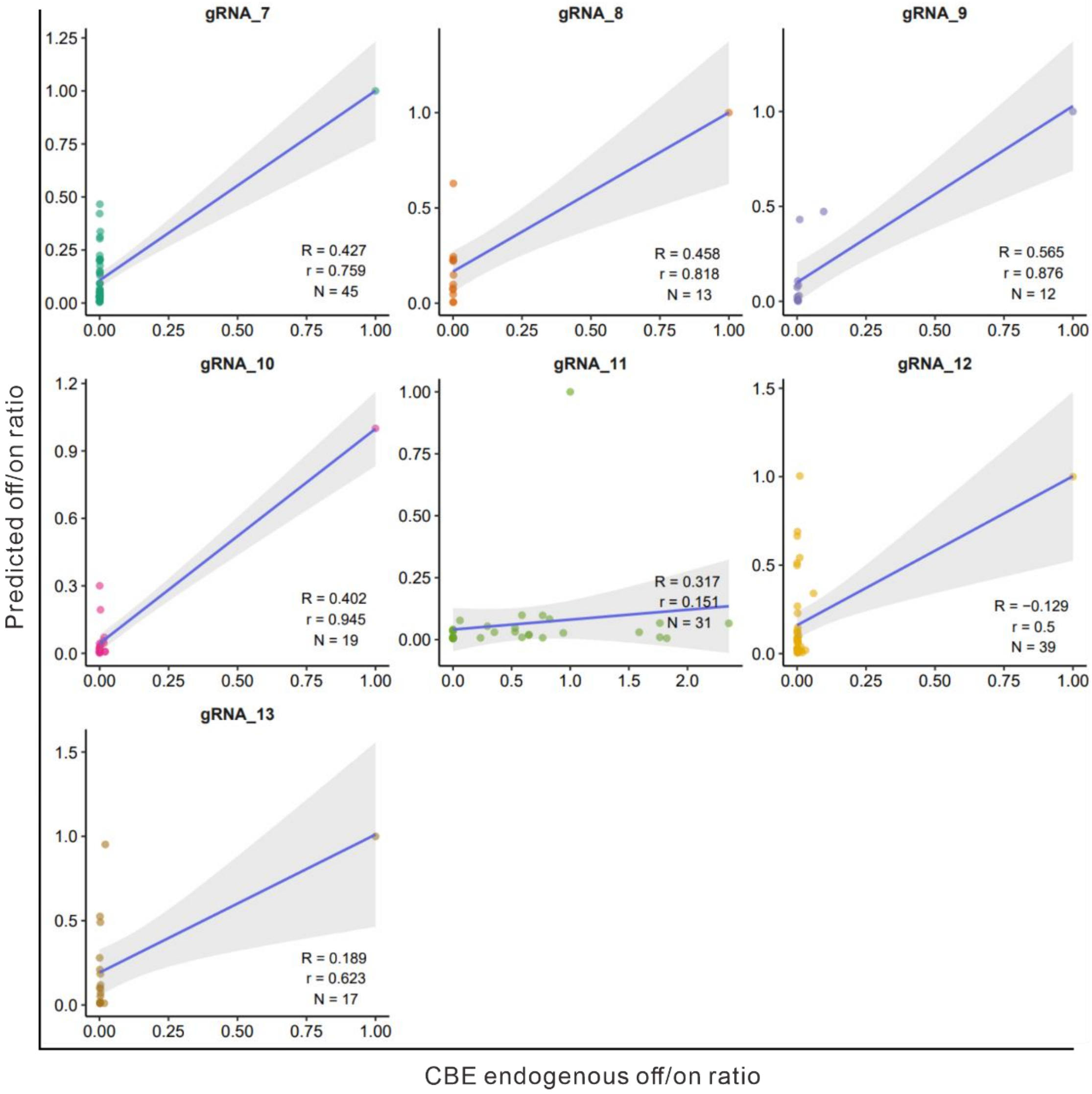


**Supplementary Figure 5. Evaluation of CBEdeepoff prediction for conversion efficiency with 7 groups of endogenous off-target datasets.**
